# Central inhibition of Stearoyl-CoA Desaturase has minimal effects on the peripheral metabolic symptoms of the 3xTg Alzheimer’s disease mouse model

**DOI:** 10.1101/2023.08.01.551291

**Authors:** Laura K. Hamilton, Paule Mbra, Sophia Mailloux, Manon Galoppin, Anne Aumont, Karl J.L. Fernandes

**Affiliations:** Research Center of the University of Montreal Hospital (CRCHUM), Montreal, Canada; Department of Neurosciences, Faculty of Medicine, Université de Montréal, Montreal, Canada; Research Center on Aging, CIUSSS de l’Estrie-CHUS, Sherbrooke, Canada; Department of Medicine, Faculty of Medicine and Health Sciences, Université de Sherbrooke, Sherbrooke, Canada

## Abstract

Evidence from genetic and epidemiological studies point to lipid metabolism defects in both the brain and periphery being at the core of Alzheimer’s disease (AD) pathogenesis. Previously, we reported that central inhibition of the rate-limiting enzyme in monounsaturated fatty acid synthesis, Stearoyl-CoA Desaturase (SCD), improves brain structure and function in the 3xTg mouse model of AD (3xTg-AD). Here, we tested whether these beneficial central effects involve recovery of peripheral metabolic defects, such as fat accumulation and glucose and insulin handling. As early as 3 months of age, 3xTg-AD mice exhibited obesity-like phenotypes including increased body weight and visceral and subcutaneous white adipose tissue deposition, as well as diabetic-like peripheral gluco-regulatory abnormalities. Intracerebral infusion of an SCD inhibitor that normalizes brain fatty acid metabolism, synapse loss and learning and memory deficits in middle-aged symptomatic 3xTg-AD mice did not affect peripheral phenotypes. This suggests that the beneficial effects of central SCD inhibition on cognitive function are not mediated by recovery of peripheral metabolic abnormalities. Given the widespread side-effects of systemically administered SCD inhibitors, these data suggest that selective inhibition of SCD in the brain may represent a clinically safer and more effective strategy for AD.

## Introduction

Alzheimer’s disease (AD) is an aging-dependent neurodegenerative disease that is most well-known for causing learning and memory loss. Indeed, much of early research on AD focused on the aggregation of amyloid-beta and tau in brain regions associated with cognition and memory, such as the hippocampus and entorhinal cortex^1^. However, numerous clinical trials have now succeeded in reducing such aggregations in AD subjects and have generally shown minor or no beneficial effects on cognition^2, 3^. With time, a broadened scope of research has revealed that AD patients harbour a plethora of other brain and body changes, many of which precede cognitive impairment, including dysregulated lipid metabolism^4–10^. Intriguingly, Dr. Alois Alzheimer himself described lipid droplets in the brains of AD patients over 100 years ago, a finding largely put aside due to the complexity of studying brain lipids^11^.

The development of new tools and technologies has made it increasingly possible to understand the role of lipid changes in health and disease. It is now clear that lipids have wide reaching functions in the body and that dysregulation of lipid metabolism is at the core of many diseases. A critical and potentially disease-initiating role of lipids in AD is supported by both genetic and environmental/lifestyle factors. For example, GWAS studies show that many AD risk genes such as APOE, TREM2, APOJ, PICALM, CLU, ABCA7 and ECHDC3 are directly involved in lipid metabolism^12^. Moreover, individuals with lipid-related metabolic syndromes such as obesity or diabetes are at increased risk of developing cognitive impairments and AD specifically^13, 14^.

In AD, both central and peripheral lipid changes have been reported in patients and animal models^4, 15–17^. Interestingly, dysregulation in levels of monounsaturated fatty acids (MUFA) have emerged in studies of AD and other neurodegenerative diseases, with remarkably beneficial effects found by lowering activity of Stearoyl-CoA desaturase (SCD), the rate limiting enzyme in MUFA synthesis^15, 18–26^. In AD specifically, we previously demonstrated that brain infusion of an SCD inhibitor into the lateral ventricle decreased local MUFA build up and resulted in reversal of the core biological deficits of AD, including cerebral inflammation, hippocampal dendritic spine loss, and learning and memory^15, 25^.

Intriguingly, SCD is also a powerful regulator of systemic metabolic fitness. Indeed, mice with whole-body knock-out of SCD family members are protected from diet-induced adiposity, exhibit increased insulin sensitivity, and altered plasma leptin and FFAs^27, 28^. These observations raised the question of whether central SCD inhibition triggers peripheral metabolic improvements and whether the cognitive improvements seen in AD mice following central SCD inhibition require amelioration of peripheral metabolic dysfunction. This seems possible, as it is well established that the brain is a crucial regulator of peripheral metabolism, including food intake and satiety, insulin secretion, hepatic glucose production, and glucose/fatty acid metabolism in adipose tissue and skeletal muscle^29, 30^. In this study we therefore tested whether altering brain lipid metabolism via central SCD inhibition could improve peripheral lipid metabolism phenotypes in AD model mice.

## Results

### Obesity-like phenotypes appear early in 3xTg-AD mice

Being overweight in midlife is linked to greater risk of developing AD^31, 32^ and brain changes in obese individuals mirror those of AD^33^.

Thus, we first aimed to characterize obesity-related defects in the slowly progressing 3xTg-AD mouse model. 3xTg-AD mice carry the human APP_Swe_, tau_P301L_, and PS1_M146V_ mutations and develop symptoms of learning and memory impairments as early as 6 months of age, prior to onset of amyloid plaques and neurofibrillary tangles^34^. We observed that young 2-3-month-old 3xTg-AD mice are already slightly heavier than WT mice (p=0.0684, unpaired t-test) and this peaked at 6-7-months (p=0.0044, unpaired t-test, Fig 1a). To determine if increased adiposity was responsible for this weight gain, we weighed white and brown fat pads of young presymptomatic 2-3-month-old mice. This showed a tendency for an increased percentage of both visceral (abdominal) (p=0.0689, unpaired t-test, Fig 1b) and subcutaneous (hindlimb) (p=0.0980, unpaired t-test, Fig 1c) white adipose tissue (WAT) in 3xTg-AD mice. We observed no change in the percentage of the thermogenic and metabolically protective brown adipose tissue (p=0.7756, unpaired t-test, Fig 1d). The increase in peripheral fat was not reflected in changes in circulating plasma levels of total free fatty acids (p=0.9386, unpaired t-test, Fig 1e) or triglycerides (p=0.6880, unpaired t-test, Fig 1f); however, the concentration of leptin, a key metabolic hormone that correlates with adiposity and is itself secreted by adipose tissue, was significantly elevated in 3xTg-AD plasma (p=0.0301, unpaired t-test, Fig 1g). This lipid accumulation shows an interesting parallel with our previous findings of increased lipid accumulation in the brains of young 2- month-old 3xTg-AD mice and in the post-mortem brains of AD subjects^15^.

**Figure 1:**
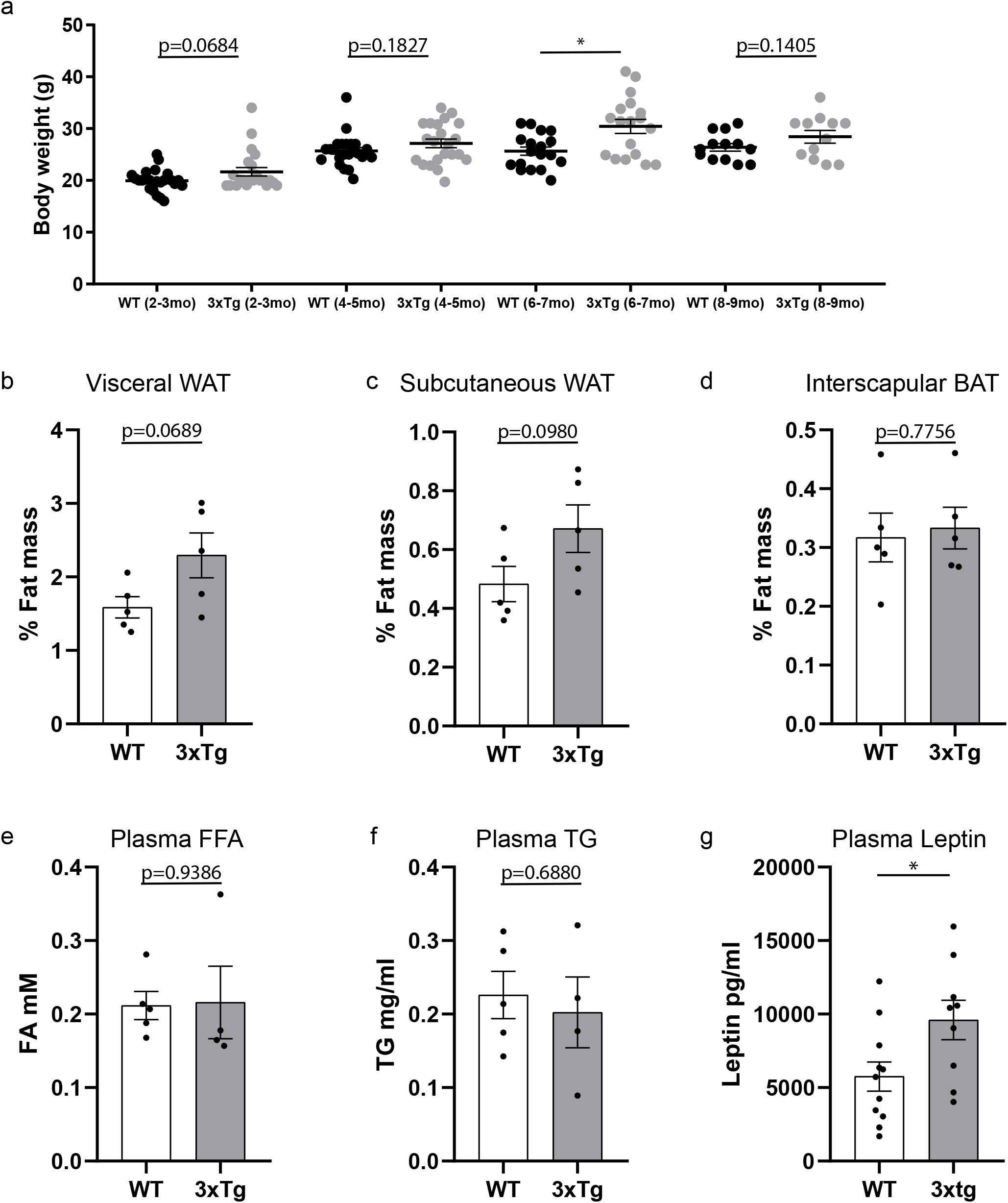
Obesity-like phenotypes appear early in --AD mice. **a** Cross sectional analysis of body weight in grams (g) between WT and 3xTg-AD mice at 2-3 months (n=22, p=0.0684), 4-5 months (n=21, p=0.1827), 6-7 months (n=18, p=0.0044), and 8-9 months (n=12-14, p=0.1405) of age. **b-d** Adipose tissue weight (g) as a percent (%) of total body weight b Visceral white (n=5, p=0.0689) c subcutaneous white (n=5, p=0.0980), d interscapular brown (n=5, p=0.7756). **e-g** Plasma concentration of e free fatty acids (n=4-5, p=0.9386), **f** triglycerides (n=4-5, p=0.6880), **g** leptin (n=9-11, p=0.0301) between 3-month-old WT and 3xTg-AD mice. Unpaired t-test. Error bars represent mean ± standard error of the mean (SEM). Significance level was set at *p* ≤ 0.05.\**p*≤0.05.

Since lipotoxicity can involve metabolic and immune dysfunction, we dissected the liver and spleen in young WT and 3xTg-AD mice to examine any gross changes in organ weight (Fig S1a,b). The liver, a central hub for lipid metabolism that regulates uptake, esterification, oxidation, and secretion of fatty acids showed no weight change between WT and 3xTg-AD mice (p=0.2074, unpaired t-test, Fig S1a). On the other hand, the spleen which plays a key role in blood filtering and regulates the secretion of proinflammatory cytokines showed significant enlargement (p=0.0128, unpaired t-test, Fig S1b). In line with this, previous studies in 3xTg-AD mice have shown alterations in the spleen including increased organ weight and T cells at older ages^35–38^.

Together, these data show that 3xTg-AD mice have obesity-like phenotypes including increased peripheral fat and leptin levels, as well as splenomegaly at presymptomatic ages.

### Diabetes-like dysfunction in insulin and glucose handling occur in young 3- month-old 3xTg-AD mice

Through its communication with other organs, adipose tissue can regulate diverse processes including insulin sensitivity and glucose metabolism^39^. Under physiological conditions, pancreatic beta cells secrete insulin post prandially to increase the storage of excess glucose in the liver to reduce blood sugar. However, in type-2 diabetes, a reduction in insulin secretion and a reduced sensitivity to insulin cause an excess of blood glucose leading to chronic hyperglycemia.

Here, we used the glucose tolerance test (GTT) as an indicator of proper peripheral insulin response/secretion. Mice were fasted 16 hours prior to the administration of a single intraperitoneal (i.p) dextrose (1g/kg) injection. Plasma glucose levels were measured prior to dextrose injection (time 0) and then every 15 minutes for 90 minutes (Fig 2a). Repeated measures (RM) analysis of variance (ANOVA) identified a significant time (F(6, 72) = 77.03, p<0.0001), strain (F(1, 12) = 22.14, p=0.0005), and interaction (F(6, 72) = 8.546, p≤0.0001), between WT and 3xTg-AD glucose levels. Post hoc multiple comparison tests showed significant strain differences at t15-75 minutes post injection (Fig 2a). In addition, 3xTg-AD mice had higher fasted blood glucose compared to WT mice, prior to dextrose injection (t0) (unpaired t-test p=0.0021, Fig 2b). The slope of the first measurement following the dextrose injection was used as an indicator of maximum tolerance to glucose and showed a significantly more pronounced glucose spike by the 3xTg-AD mice (unpaired t-test p=0.0037, Fig 2c). Similarly, the area of the curve (AOC)^40^ showed a significant impairment in clearance of blood glucose, suggesting reduced glucose tolerance by the 3xTg-AD mice (unpaired t-test, p=0.0024, Fig 2d). Together, the GTT test showed dysregulation of glucose handling in young 3xTg-AD mice.

**Figure 2:**
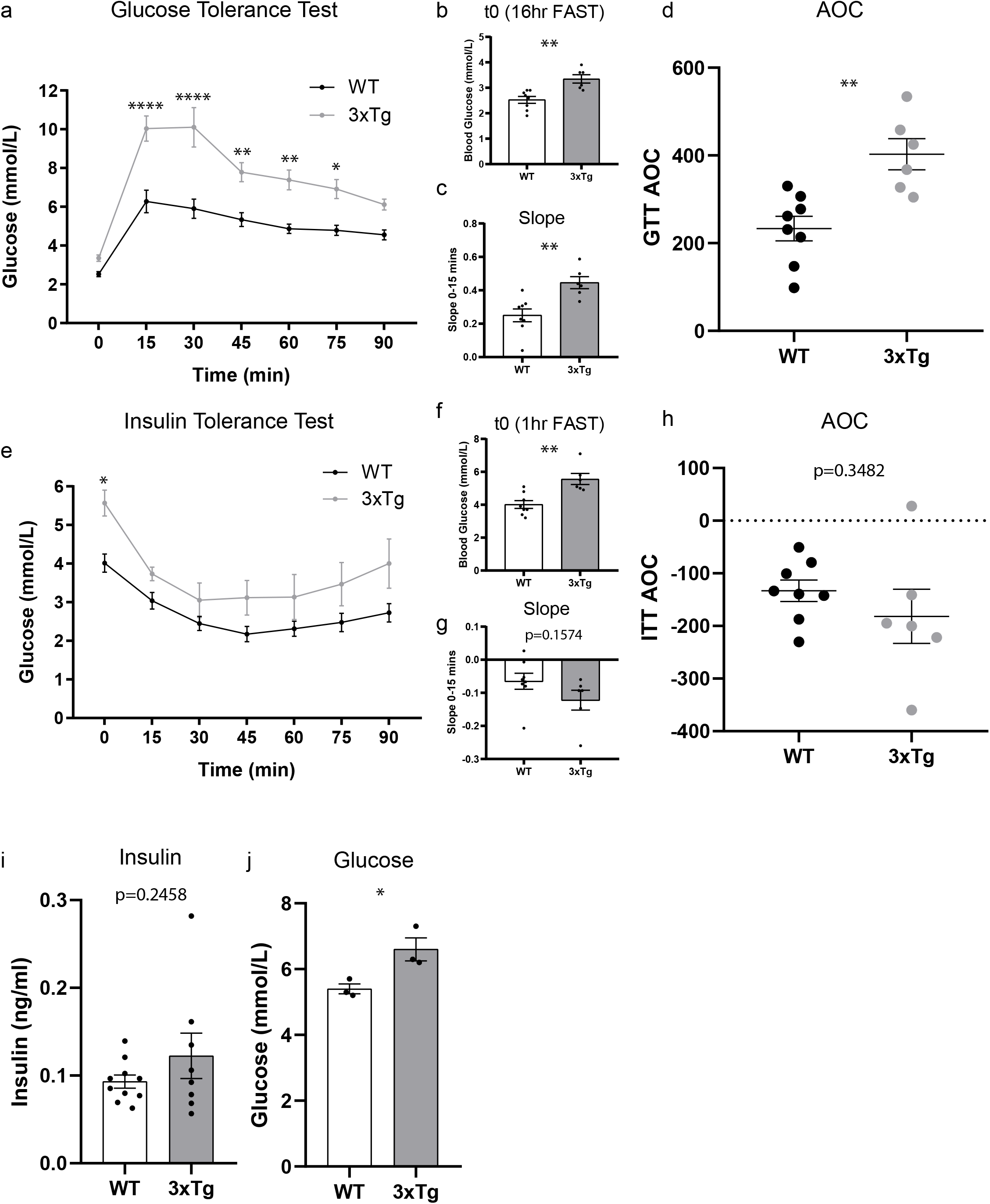
Diabetes-like dysfunction in insulin and glucose handling occur in young 3-month-old 3xTg-AD mice. **a**-**d** Glucose tolerance test (GTT), 16-hour fasted mice were given an intraperitoneal injection of dextrose (1 mg/g body weight). Glucose concentration (mmol/L) was measured from blood sampled from the tail vein at the times indicated (n=6-8 mice/group). **a** Repeated measures (RM) analysis of variance (ANOVA) identified a significant time (F(6, 72) = 77.03, p≤0.0001), strain (F(1, 12) = 22.14, p=0.0005), and interaction (F(6, 72) = 8.546, p≤0.0001) between WT and 3xTg-AD glucose levels. Post hoc multiple comparison analysis showed significant differences at t15-75 minutes post injection. **b** blood glucose levels between WT and 3xTg-AD mice prior to injection of glucose (p=0.0021). **c** Slope of the first 15 minutes after glucose injection (p=0.0037). **d** Area of the curve of the GTT (p=0.0024). **e**-**h** Insulin tolerance test (ITT), 16-hour fasted mice were given an intraperitoneal injection of insulin (0.5U/g body weight). Glucose concentration (mmol/L) was measured from blood sampled from the tail vein at the times indicated. **e** RM-ANOVA identified significant time (F(1.984, 23.81) = 18.89, p≤0.0001) and strain effects (F(1, 12) = 7.555, p=0.0176) between WT and 3xTg-AD glucose levels, but no significant interaction. Post hoc multiple comparison analysis showed significant differences at t15 minutes post injection. **f** Blood glucose levels between WT and 3xTg-AD mice prior to injection of insulin (p=0.0021). **g** Slope of the first 15 minutes after insulin injection between WT and 3xTg mice (p=0.1574). **h** Area of the curve of the ITT between WT and 3xTg-AD mice (p=0.3482). **i-j** unfasted plasma i insulin concentration (ng/ml) (n=8-10, p=0.2458), **j** glucose concentration (mmol/L) (n=3, p=0.0314), between WT and 3xTg-AD mice. Unpaired t-test. Error bars represent mean ± standard error of the mean (SEM). Significance level was set at *p* ≤ 0.05. *≤0.05, **≤0.05, ****≤0.0001.

Insulin suppresses glucose release from the liver and promotes glucose uptake by muscle. The degree to which blood glucose levels fall in response to insulin administration is indicative of insulin sensitivity. To assess if the 3xTg-AD mice exhibit signs of insulin resistance/insensitivity, we employed the insulin tolerance test (ITT). Mice were fasted 1 hour prior to the administration of a single i.p insulin (0.5U/kg) injection. Plasma glucose levels were taken prior to insulin injection for baseline (time 0) assessment and every 15 minutes for 90 minutes (Fig 2e). RM-ANOVA identified a significant time (F(1.984, 23.81) = 18.89, p<0.0001), strain (F(1, 12) = 7.555, p=0.0176), and no significant interaction (F(6, 72) = 0.9370, p=0.4741) between WT and 3xTg-AD glucose levels. Post hoc multiple comparison tests showed significant strain differences at t15 minutes post injection (Fig 2e). In addition, the 3xTg-AD mice showed higher baseline glucose compared to WT mice (unpaired t-test, p=0.0021, Fig 2f). The slope of the first measurement following the insulin injection showed a trend towards a sharper response by the 3xTg-AD mice (unpaired t-test, p=0.1574, Fig 2g). The area of the curve (AOC) showed no significant difference (unpaired t-test, p=0.3482, Fig 2h). Unfasted insulin was not significantly changed (unpaired t-test, p=0.2458, Fig 2i) while unfasted glucose was significantly elevated in 3xTg-AD mice (unpaired t-test, p=0.0314, Fig 2j). Moreover, the weight of the insulin-secreting pancreas was unchanged at these ages (Fig S1c). In sum, the ITT results showed dysregulation of insulin handling in young 3xTg-AD mice.

Together, these data show that 3xTg-AD mice have increased fat and gluco- regulatory abnormalities at presymptomatic ages making them a good model to study the peripheral effects of the lipid lowering drug SCD.

### 1-month central SCDi does not alter peripheral fat accumulation

Previous studies have shown increased gene expression of Scd1 in 3xTg-AD mice and SCD/SCD5 in AD patients, along with changes in their fatty acid substrates and products^10, 15, 24, 25, 41^. We used intracerebroventricular infusion of an SCD inhibitor (SCDi) and found marked changes in overall hippocampal gene expression that results in widespread cellular benefits, including a decrease in microglia activation, rescue of dendritic spines and structure and a functional rescue of learning and memory deficits in symptomatic 3xTg-AD mice^15, 25^. SCDs are central lipogenic enzymes throughout the body, catalyzing the desaturation of the saturated fatty acids (SFA), palmitate and stearate, into the MUFAs, palmitoleate and oleate, respectively. As mentioned previously, whole body knock-out, including the brain, of Scd1 in mice improves metabolic fitness including reduced adiposity, plasma leptin and FFAs^27, 28^. Given the importance of the CNS in regulation of peripheral metabolism, we asked whether the beneficial effects of central inhibition of SCD involve and might be mediated by improvements in peripheral metabolism.

To assess if central SCD inhibition improves peripheral metabolic features of AD, we employed the same protocol as previously^25^, infusing SCDi ICV for 1-month in symptomatic 9-month-old WT and 3xTg-AD mice (Fig 3a). Comparison of body weight at the end of the 1-month study (Fig 3b) showed a significant strain effect with 3xTg-AD mice weighing more than WT mice (F(1,48) = 21.26, p<0.0001) but no drug effect or interaction. Post hoc analysis showed a significant difference between vehicle treated WT-V/3xTg-V (p=0.0456) and WT-S/3xTg-S (p=0.0023) and no significant difference between 3xTg-V/3xTg-S (p=0.4294), 2-way ANOVA Tukey’s post hoc. Body composition analysis by EchoMRI showed a significant strain effect with 3xTg-AD mice having more fat than WT mice (F(1,26) = 29.19, p<0.0001) but no drug effect or interaction. Post hoc analysis showed a significant difference between vehicle treated WT-V/3xTg-V (p=0.0040) and WT-S/3xTg-S (p=0.0038) and no significant difference between 3xTg-V/3xTg-S (p=0.9418), 2-way ANOVA Tukey’s post hoc. Lean mass was unchanged between strains, drug, and showed no interaction. While fat as a percent of total body weight showed a significant strain effect (F(1,26) = 37.36, p p<0.0001) but no drug effect or interaction. Post hoc analysis showed a significant difference between vehicle treated WT-V/3xTg-V (p=0.0012) and WT-S/3xTg-S (p=0.0010) and no significant difference between 3xTg-V/3xTg-S (p=0.8903), 2-way ANOVA Tukey’s post hoc.

**Figure 3:**
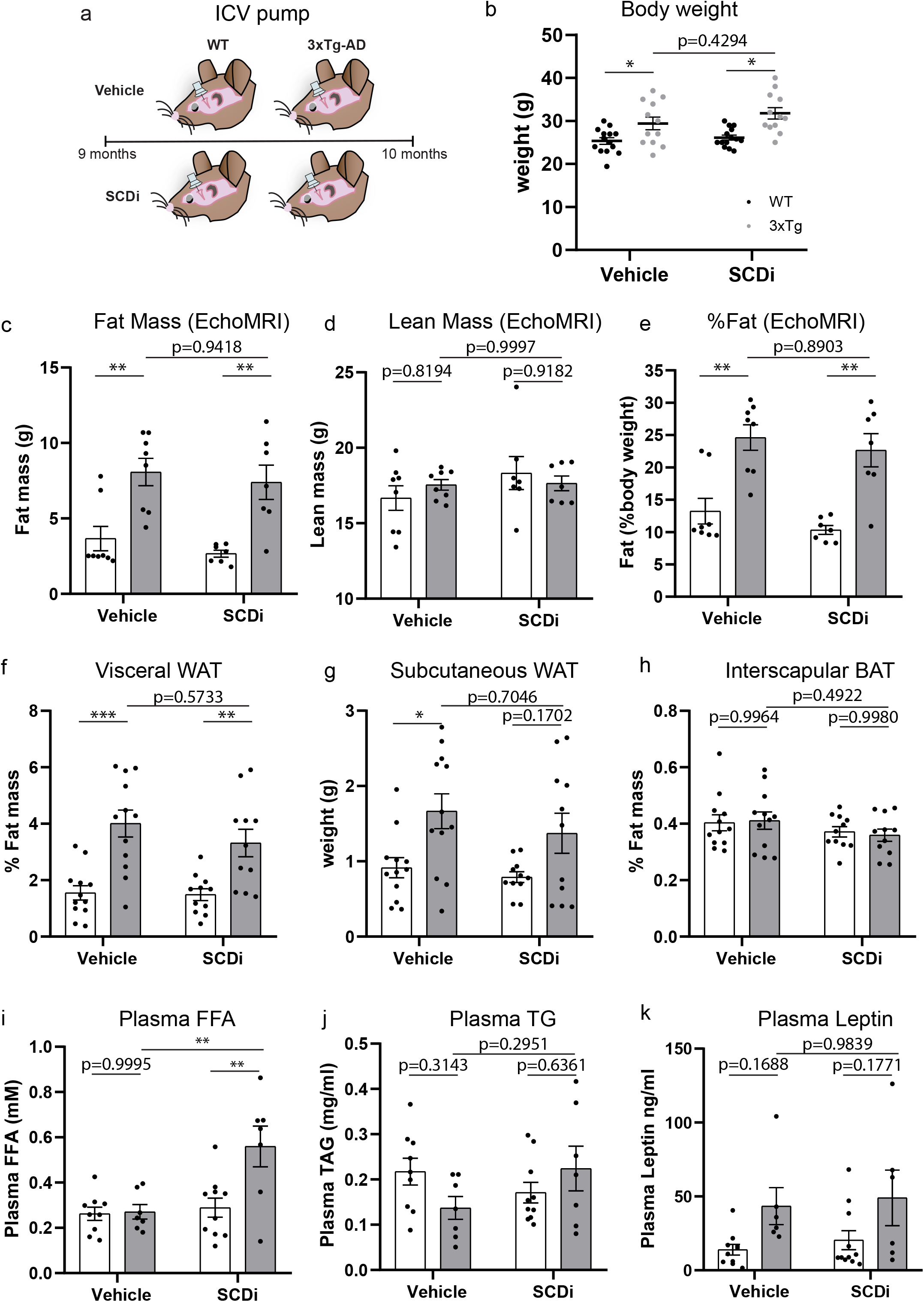
1-month central SCDi does not alter peripheral fat accumulation. **a** Schematic of experimental groups, 9-month-old WT and 3xTg-AD implanted with intracerebroventricular osmotic pumps containing vehicle (V) or SCD inhibitor (S) for 1- month. **b**, Body weight in grams (g) (n=12-14 animals/group) at sacrifice showed a significant strain effect with 3xTg-AD mice weighing more than WT mice (F(1,48) = 21.26, p<0.0001) but no drug effect or interaction. Tukey’s post hoc analysis showed a significant difference between vehicle treated WT-V/3xTg-V (p=0.0456) and WT- S/3xTg-S (p=0.0023) and no significant treatment effect 3xTg-V/3xTg-S (p=0.4294). **c-e**, Body composition by EchoMRI (n=7-8 animals/group). **c** 2-way ANOVA showed a significant strain effect with 3xTg-AD mice having more fat than WT mice (F(1,26) = 29.19, p<0.0001) but no drug effect or interaction. Tukey’s post hoc analysis showed a significant difference between vehicle WT-V/3xTg-V (p=0.0040) or drug treated WT- S/3xTg-S (p=0.0038) and no significant difference between 3xTg-V/3xTg-S (p=0.9418). **d** lean mass was unchanged between strains, drug, and showed no interaction. e fat mass as a percentage of body weight showed a significant strain effect (F(1,26)=37.36, p<0.0001) but no drug effect or interaction. Tukey’s post hoc analysis showed a significant difference between vehicle treated WT-V/3xTg-V (p=0.0012) and WT- S/3xTg-S (p=0.0010) and no significant difference between 3xTg-V/3xTg-S (p=0.8903). **f-h** Adipose tissue weight in grams (g) as a percent (%) of total body weight (n=11-12 animals/group). f visceral white g, subcutaneous white, h, interscapular brown. A significant strain effect on f visceral (F(1,42)=31.87, p<0.0001), post hoc analysis showing a significant difference between vehicle treated WT-V/3xTg-V (p=0.0002) and WT-S/3xTg-S (p=0.0095) and no significant difference between 3xTg-V/3xTg-S (p=0.5733), 2-way ANOVA Tukey’s post hoc. **g** subcutaneous WAT also showed a significant strain effect (F(1,42)=12.04, p=0.0012) but no drug or interaction. Post hoc analysis showed a significant difference between vehicle treated WT-V/3xTg-V (p=0.0348) and WT-S/3xTg-S (p=0.1702) and no significant difference between 3xTg- V/3xTg-S (p=0.7046), 2-way ANOVA Tukey’s post hoc. Moreover, we did not detect any changes in BAT % in any group **h**. **i-k** Plasma concentration of i free fatty acids (n=7-10 animals/group), j triglycerides (n=7-10 animals/group), k leptin between WT and 3xTg-AD mice (n=6-11 animals/group). No effect was observed between vehicle treated WT and 3xTg-AD mice (p=0.9995, 2-way ANOVA Tukey’s post hoc), however, plasma free fatty acids were elevated in 3xTg-AD mice treated with SCDi compared to Vehicle (p=0.0043, 2-way ANOVA Tukey’s post hoc). **j** no significant strain or drug effect on total triglycerides was detected, 2-way ANOVA. **k** levels of leptin showed a significant strain effect (F(1,29) = 8.929, p=0.0057) but no drug effect or interaction. Post hoc analysis showed no significant differences between any groups, 2-way ANOVA Tukey’s post hoc. Error bars represent mean ± standard error of the mean (SEM). Significance level was set at *p* ≤ 0.05.*p≤0.05, **p≤0.01, ***p≤0.001.

Since young 2–3-month-old mice had an increase in WAT (Fig 1) we again dissected fat pads from visceral (abdominal) WAT (Fig 3f), subcutaneous (hind limb) WAT (Fig 3g) and interscapular BAT (Fig 3h). We found a significant strain effect on visceral WAT %(F(1,42)=31.87, p<0.0001), with post hoc analysis showing a significant difference between vehicle treated WT-V/3xTg-V (p=0.0002) and WT-S/3xTg-S (p=0.0095) and no significant drug effect 3xTg-V/3xTg-S (p=0.5733), 2-way ANOVA Tukey’s post hoc. Subcutaneous WAT % also showed a significant strain effect (F(1,42)=12.04, p=0.0012) but no drug or interaction. Post hoc analysis showed a significant difference between vehicle treated WT-V/3xTg-V (p=0.0348) and WT- S/3xTg-S (p=0.1702) and no significant drug effect 3xTg-V/3xTg-S (p=0.7046), 2-way ANOVA Tukey’s post hoc. Moreover, we did not detect any changes in BAT % in any group at this older symptomatic age (Fig 3h).

To determine if plasma metabolites were altered by ICV-SCDi, we measured plasma free fatty acids, triglycerides, and leptin levels. Intriguingly, although there was no baseline strain difference in plasma free fatty acid levels between WT and 3xTg-AD mice, 3xTg-AD mice treated with ICV SCDi showed a significant increase (3xTg- V/3xTg-SCDi, p=0.0043, 2-way ANOVA, Tukey’s post hoc), suggestive of fatty acid clearance from the brain into the blood (Fig 3i). No significant strain or drug effect on total triglycerides was detected (Fig. 3j, 2-way ANOVA), however, levels of the adipose signalling hormone leptin showed a significant strain effect (F(1,29) = 8.929, p=0.0057) but no drug effect or interaction. Post hoc analysis showed no significant differences between any groups, 2-way ANOVA Tukey’s post hoc.

In the mouse, Scd1 is ubiquitously expressed and shows highest expression in highly metabolic tissues. To determine if ICV-SCD altered peripheral organ weight we measured the weight of the liver and spleen. No significant treatment or strain changes were observed on liver weight (Fig S2a). As at younger ages, the spleen (Fig S2b) showed a significant strain effect (F(1,29) =8.929, p<0.0001). Post hoc analysis showed a significant difference between vehicle WT-V/3xTg-V (p=0.0157) and SCDi treated WT- S/3xTg-S (p=0.0009) but no significant drug effect, 3xTg-V/3xTg-S (p=0.5725), 2-way ANOVA Tukey’s post hoc.

Together, these data demonstrate that ICV SCDi for one month does not alter the gross peripheral fat accumulation and associated circulating leptin levels seen in the 3xTg-AD mice.

### 1-month central SCDi does not improve insulin and glucose handling

Since young 3xTg-AD mice exhibited impaired glucose regulation (Fig. 2) and glucose sensitivity is improved in whole-body Scd1-KO mice, we next assessed the impact of central SCDi infusion on glucose and insulin tolerance.

The GTT test (Fig 4a) showed significant group (F(3, 25) =3.078, p=0.0458) and time effects (F(3.865,96.62)=36.24, p<0.0001) with no interaction by 2-way RM-ANOVA. Indeed, resting glucose levels were significantly higher in 3xTg-AD mice compared to WT (F(1,25)=13.45, p=0.0012). Tukey’s post hoc analysis showed a trend towards higher resting glucose levels in 3xTg-V mice (p=0.0863), but no drug effect or interaction was observed (Fig 4b). The slope of the first measurement following the dextrose injection was used as an indication of the tolerance to glucose and showed no strain or drug effect, 2-way ANOVA (Fig 4c). Similarly, the area of the curve (AOC) showed no strain or treatment effect on the response to glucose, 2-way ANOVA (Fig 4d).

**Figure 4:**
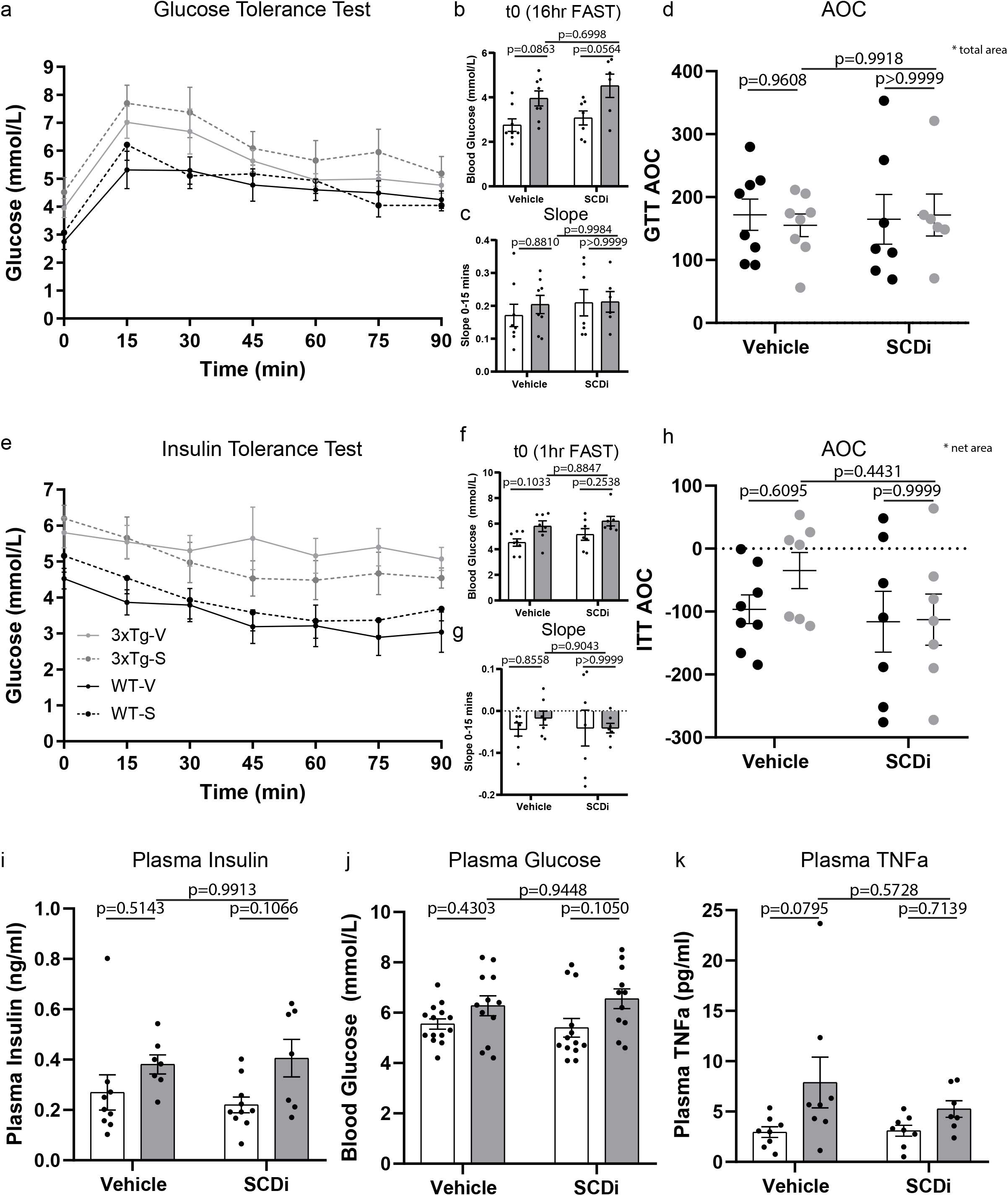
1-month central SCDi does not improve insulin and glucose handling. **a**-**d** Glucose tolerance test (GTT), 16-hour fasted mice were given an intraperitoneal injection of dextrose (1 mg/g body weight, n=7-8 animals/group). **a** Glucose concentration (mmol/L) was measured from blood sampled from the tail vein at the times indicated. 2-way RM-ANOVA showed a significant group (F(3, 25) =3.078, p=0.0458) and time effects (F(3.865,96.62)=36.24, p<0.0001) with no interaction. **b** Blood glucose levels between WT and 3xTg-AD mice prior to injection of glucose (time 0) were significantly higher in 3xTg-AD mice compared to WT (F(1,25)=13.45, p=0.0012). Tukey’s post hoc analysis showed a trend towards higher resting glucose levels in 3xTg-V mice (p=0.0863), but no drug effect or interaction. **c** Slope of the first 15 minutes after glucose injection between WT and 3xTg-AD mice showed no strain, drug effect, or interaction by 2-way ANOVA. **d** Area of the curve of the GTT between WT and 3xTg-AD mice by 2-way ANOVA showed no strain or treatment effect. **e**-**h** Insulin tolerance test (ITT), 1-hour fasted mice were given an intraperitoneal injection of insulin (0.5U/g body weight, n=7-8 animals/group). **e** glucose concentration (mmol/L) was measured from blood sampled from the tail vein at the times indicated. The ITT test showed significant group (F(3,25)=4.675, p=0.0100) and time effects (F(3.315, 82.89)=14.16, p<0.0001). **f** Blood glucose levels between WT and 3xTg-AD mice prior to injection of insulin (time 0) showed higher baseline glucose in 3xTg-AD mice compared to WT (2-way ANOVA, (F(1,25)=9.200, p=0.0056) but no drug effect or interaction. **g** Slope of the first 15 minutes after insulin injection between WT and 3xTg- AD mice showed no strain or treatment effect, 2-way ANOVA. **h** Area of the curve of the ITT between WT and 3xTg-AD mice showed no strain or treatment effect, 2-way ANOVA. **i-k** unfasted plasma i insulin concentration (n=7-10 animals/group) showed a strain effects (F(1,46)=7.513, p=0.0087), **j** glucose concentration (n=11-14 animals/group) showed a strain effect (F(1,29)=7.020, p=0.0129) **k** TNFa (n=7-9 animals/group) showed a strain effect (F(1,27)=6.326, p=0.0182), by 2-way ANOVA. No significant drug effect or interaction was observed between WT and 3xTg-AD mice. Error bars represent mean ± standard error of the mean (SEM). Significance level was set at *p* ≤ 0.05.

To evaluate signs of insulin resistance/insensitivity we performed the ITT test (Fig 4e). The ITT test showed a significant group (F(3,25)=4.675, p=0.0100) and time effects (F(3.315, 82.89)=14.16, p<0.0001). The 3xTg-AD mice showed higher baseline glucose compared to WT mice (2-way ANOVA, (F(1,25)=9.200, p=0.0056) but no drug effect or interaction was observed (Fig 4f). The slope of the first measurement following the insulin injection was used to infer the response to insulin and showed no strain or treatment effect, 2-way ANOVA (Fig 4g). The area of the curve (AOC) also showed no strain or treatment effect, 2-way ANOVA (Fig 4h). To assess the unfasted resting glucose and insulin levels we measured resting plasma glucose (Fig 4i) and insulin (Fig 4j) levels and found significant strain effects (F(1,46)=7.513, p=0.0087), (F(1,29)=7.020, p=0.0129) respectively, 2-way ANOVA, but no significant effect of treatment or interaction. We also detected no weight difference in insulin secreting pancreas at these ages (Fig S2c).

Increased adiposity causes inflammation and increased cytokine production that can impact both the brain and periphery^42^. We have previously shown in LPS-treated mouse microglia, that SCDi or Scd1-KO dampens Tumor necrosis factor alpha (TNFa)^25^. TNFa is a cytokine and adipose-tissue-associated adipokine known to contribute to insulin resistance associated to obesity^43^. Here, plasma TNFa levels showed a strain dependent increase in 3xTg-AD mice (F(1,27)=6.326, p=0.0182), 2-way ANOVA) however, no significant drug effect or interaction was observed (Fig 4k). Thus, these data show that the significantly higher peripheral fat accumulation and cytokines in 3xTg-AD mice are not significantly modulated following 1-month central administration of SCDi.

Collectively, these findings show that 3xTg-AD mice have increased resting and fasted glucose levels, that their response to insulin and glucose challenge is moderately affected in mid-life, and that these metabolic features are not modulated by 1-month central administration of SCDi.

## Discussion

This study assessed the ability of central SCD inhibition to regulate peripheral metabolism in the 3xTg-AD mouse model. Using a genetic model of AD where dementia-causing mutations lead to cognitive impairment in mid-life, we confirmed that 3xTg-AD mice exhibit gross metabolic deficits that precede these cognitive symptoms. Moreover, we show that while 3xTg-AD mice have early and mid-life metabolic changes, including increased peripheral fat accumulation and circulating leptin as well as impaired resting glucose and glucose tolerance, 1-month ICV infusion of SCDi in mid-life does not normalize these parameters.

The ICV infusion paradigm for SCDi used here results in remarkable improvements in brain structure and function of 3xTg-AD mice, including a recovery of learning and memory in mid-life^25^. Knowing that these mice also have peripheral metabolic deficits, we wondered if the benefits may be due in part to improved metabolism in the periphery. Our present results reveal that 1-month ICV SCDi does not significantly affect the aspects of peripheral metabolism measured here, indicating that improvements of these peripheral metabolic features are not needed for the benefits of central SCDi treatment on the brain. This is important as it suggests that targeting CNS MUFA metabolism may be sufficient to have cellular and cognitive improvements in AD. Intriguingly, since SCDi-induced increases in levels of plasma free fatty acids were detected, it is conceivable that longer term CNS treatment with SCDi may eventually impact peripheral metabolism.

### AD and peripheral metabolism

AD patients exhibit significant metabolic symptoms including body weight changes, dyshomeostasis of central and peripheral lipids, diminished brain glucose uptake, and Type 2 diabetes. Although these alterations occur prior to the initial mental decline and progressively worsen as the disease advances, it is difficult to establish whether they are AD prodromes, risk factors, or both. Our data suggests that expression of dementia-causing genes (APP, PS1, Tau) is sufficient to cause peripheral metabolic deficits that are reminiscent of obesity and diabetes and that occur in the absence of changes in dietary or environmental factors. Indeed, other groups have also reported changes in peripheral parameters in human familial AD patients and genetic AD animal models^5^. These observations are in line with the idea that treating the CNS sufficiently early could interrupt the cascade that leads to neurodegeneration and peripheral metabolic syndromes.

### Pleiotropic effects of SCD inhibition in peripheral metabolism

Studies in transgenic mouse models have demonstrated an essential role of Scd1 in regulating cellular processes including lipid synthesis and oxidation, thermogenesis, hormonal signaling, and inflammation. Scd1 was initially identified in the ‘asebia’ mouse that has a naturally occurring mutation in the *Scd1* gene and exhibits alopecia, sebocyte hypoplasia, and resistance to leptin deficiency induced obesity^44^. Indeed, Scd1 itself is a known target of leptin and many of leptin’s metabolic effects are mediated through its effects on Scd1^45–47^. Subsequently, whole-body Scd1 knock-out mice were engineered and found to be resistant to high-fat diet induced obesity, with reduced hepatic as well as circulating triglycerides and cholesterol esters, dry eye, alopecia, dermatitis, and increased skin barrier permeability^28, 48^. These intriguing results spawned multiple tissue-specific Scd1 deletion studies. For instance, liver-specific Scd1 deficiency protects mice from carbohydrate-induced *de novo* lipogenesis^49^. Adipose- specific deficiency results in decreased inflammation in adipocytes and increased insulin-independent glucose uptake^50^. Interestingly, the obesity-resistant lean metabolic phenotype of global Scd1 deficiency was recapitulated not by liver or adipose-specific Scd1 deletion but by deletion in the skin^51^. Deficiency of Scd1 in the skin resulted in significant increases in energy expenditure and protection from diet-induced obesity, hepatic steatosis, and glucose intolerance. These mice also had an inability to retain heat due to significant alopecia and poor skin integrity.

Together, this suggests that SCD has far reaching effects on central and peripheral metabolism, increasing the risk of unwanted secondary effects from systemically administered inhibitors.

### SCDi as a therapeutic target for neurodegenerative diseases

This study in combination with our prior findings^15, 25^ shows that CNS SCDi is sufficient to have major beneficial effects on core AD symptoms including cognition and inflammation. Indeed, Scd inhibition is showing beneficial effects in multiple neurodegenerative disease models including Multiple sclerosis (MS), Parkinson’s disease (PD), and AD^15, 18–23, 25, 26^. However, the studies in MS and PD have used oral SCDi, making it unclear if the benefits come from the CNS or periphery. A clinical trial using oral administration of an SCDi in PD patients was launched by Yumanity therapeutics in 2021, and the results of this trial will provide vital information on the feasibility of systemically administered SCDi to treat neurodegenerative diseases.

When thinking of translating SCDi into a clinical setting, it is essential to consider the most specific target and most effective mode of administration to minimize off-target and unwanted side-effects. Our data suggests that administration of SCDi into the CNS will be sufficient to have major beneficial effects on AD and potentially other neurodegenerative diseases. Future studies using CNS-specific SCDi in other neurodegenerative diseases will be essential to determine if this is the case. In this regard, delivery systems that are better able to selectively target a genes and locations of interest, such as antisense oligonucleotides and nasal sprays, are already used in the clinic for other indications; our data supports developing such modes of administration for SCD inhibition in AD as well.

## Methods

Experiments were approved by the *Institutional Animal Care Committee* of the Centre de Recherche du Centre Hospitalier de l’Université de Montréal (CRCHUM) following the Canadian Council of Animal care guidelines.

### 3xTg-AD mice and strain controls

3xTg-AD mice (Jackson Laboratory MMRRC stock #: 034830) and their WT strain controls (B6129SF2/J, Jackson Laboratory stock #: 101045) were purchased from Jax mice. Briefly, 3xTg-AD mice were originally derived by Oddo and colleagues by co-microinjecting two independent transgenes encoding human APP_Swe_ and the human tau_P301L_ (both under control of the neuron-specific mouse Thy1.2 regulatory element) into single-cell embryos harvested from homozygous mutant PS1_M146V_ knock in (PS1- KI) mice^34^. Wildtype mice are the PS1-knock-in background strain (C57BL/6J x 129S1^34^. All mice were bred in-house, maintained in identical housing conditions (22°C, 50% humidity, 12h Light/Dark cycle OFF 10am ON 10pm) and given free access to a standard rodent diet (#2018, Harlan Teklad) and water. 3xTg-AD mice undergo a progressive increase in amyloid beta peptide deposition, with intracellular immunoreactivity being detected in some brain regions as early as 3-4 months. Synaptic transmission and long-term potentiation are demonstrably impaired in mice 6 months of age. Evidence of gliosis and inflammation is present by at least 7 months^52^. Between 12-15 months aggregates of conformationally altered and hyperphosphorylated tau are detected in the hippocampus^34, 53^. Female mice were used for all experiments and sex and age matched.

### Metabolic profiling

#### Insulin tolerance test (ITT)/Glucose tolerance test (GTT)

For the glucose tolerance test (GTT), mice were fasted 16 hours (5pm-9am) followed by i.p injection of dextrose (1 g/kg of body weight). For insulin tolerance tests (ITT), mice were fasted for 1 hour (12-1pm) prior to intraperitoneal (i.p.) injection of insulin (0.5U/kg). For both GTT and ITT, glucose concentration (mmol/L) was measured using blood glucose strips and corresponding glucometer (FreeStyle Lite, Abbott) from blood sampled from the tail vein at 0, 15-, 30-, 45-, 60-, and 90-minutes post injection. Slope of the first 15 minutes was achieved by linear regression fitting in GraphPad Prism Version 8 (GraphPad Software, Inc). Area of the curve (AOC) was calculated by using the pre-injection glucose concentration (time 0) of each mouse as baseline and including peaks below zero for ITT and only positive peaks for GTT.

#### Body composition

Fat and lean mass was measured using an EchoMRI-100 body composition analyzer (version 2008.01.18, EchoMRI LLC) at the rodent cardiovascular core facility of the CRCHUM. % fat mass and % lean mass was obtained by dividing by total body weight (g).

#### Organ weights

Visceral (abdominal) white adipose tissue, subcutaneous (hind limb) white adipose tissue and brown (interscapular) adipose tissue were dissected and weighed. % organ weight was calculated by dividing the organ weight by the total body weight (g). Whole liver, spleen and pancreas were dissected and weighed.

#### Blood collection

Blood was collected from the posterior *vena cava* at the time of sacrifice. Samples were transferred to a microtainer containing K_2_EDTA (BD Biosciences) inverted 20 times, and the plasma extracted after centrifuging at 1000 g for 20 minutes at 4°C. All samples were stored at −80°C until use.

#### Insulin, Leptin and TNFa plasma measurements

AlphaLISA assay kits (#AL204 for insulin, AL521 for Leptin and AL505 for TNFa) from PerkinElmer (Waltham, MA) and an Envision 2104 plate reader from the same supplier were used to assess plasma levels of the corresponding analyte using white 96-well microplates. A standard curve was run on each plate containing samples. Omnibead calibrators (PerkinElmer) were utilized to ensure proper operation of the assay and instrument. The analyses were carried out at the Cellular Physiology core facility of the CHUM Research Center.

#### Free fatty acids and triglycerides measurements

Plasma Triglycerides (TG) were evaluated using a commercial kit (GPO Trinder, Sigma-Aldrich, Saint Louis, USA) in duplicate. Calculations were performed to quantify triacylglycerol levels using triolein equivalent as per manufacturer recommendations. Plasma FFA were assessed using the NEFA C kit in duplicate (Wako Chemicals, Neuss, Germany) according to the manufacturer’s protocol. The analyses were carried out at the Cellular Physiology core facility of the CHUM Research Center.

### SCD inhibitor

SCD inhibitor used in this study: ab142089 (Abcam, C20H22ClN3O3, 4-(2- Chlorophenoxy)-*N*-[3 [(methylamino)carbonyl]phenyl]-1-piperidinecarboxamide, M.W. 387.87).

### In vivo surgical procedures

#### Intracerebroventricular (ICV) osmotic pumps

For ICV infusions of SCDi or vehicle, mice were locally injected with buprivacaine (Hospira) and operated under isoflurane anesthesia (Baxter). Brain cannulae attached to Alzet osmotic pumps were stereotaxically implanted at 0.0mm antero-posterior and 0.9mm lateral to Bregma and the pumps placed under the back skin according to manufacturer’s instructions. The 28-day Alzet osmotic pumps used in these experiments (0.11 μl/h infusion rate, model 1004; Durect) were primed for 48 hrs and began pumping immediately when implanted. The SCD inhibitor was dissolved in DMSO (Sigma-Aldrich) and infused at a final concentration of 80 µM in sterile aCSF (148mM NaCl, 3mM KCl, 1.7mM MgCl_2_, 1.4mM CaCl_2_, 1.5mM Na_2_PO4, 0.1mM NaH_2_PO_4_). Vehicle pumps contained the same volumes as the SCDi pumps (0.8% DMSO/aCSF).

### Statistical Analyses

All statistical analyses were performed using GraphPad Prism, Version 8 (GraphPad Software, Inc). When comparing one independent variable a t-test was used, otherwise a 2x2 experimental design (strain and treatment) was chosen and therefore analyzed by 2-way ANOVA followed by multiple comparison by Tukey’s post-hoc (all group comparisons) tests as indicated in the legends. Error bars represent mean ± standard error of the mean (SEM). Significance level was set at *p* ≤ 0.05 as indicated in the figure legends. *p≤0.05, **p≤0.01, ***p≤0.001, ****p≤0.0001.

## Acknowledgments

This study is funded by of Health Research (CIHR), the Alzheimer Society of Canada, the Natural Sciences and Engineering Research Council (NSERC) and Fondation du CHUM (Renée Claude charitable fund for Alzheimer’s research). L.K.H was supported by a Spark Award fellowship from the Canadian Alzheimer Society Research Program (ASRP).

## Author Contributions

Conceptualization, L.K.H., K.J.L.F.; Methodology, L.K.H.; Investigation, L.K.H.; P.M., S.M., M.G., A.A.; Formal analysis, L.K.H.; Writing – Original Draft, L.K.H.; Writing – Review & Editing, L.K.H., K.J.L.F.; Funding Acquisition, K.J.L.F.; Resources, K.J.L.F.; Supervision, K.J.L.F.

## Declaration of interests

The authors declare no competing interests.

## Supplemental Material

### Supplemental Figure legends

**Figure S1: Peripheral organ weight in 3-month-old WT and 3xTg-AD mice**

**a**-**c** Organ weight in grams (g), n=5 animals/group. **a** total liver weight showed no change between WT and 3xTg-AD mice (p=0.2074, unpaired t-test). **b** total spleen weight showed significant enlargement in 3xTg-AD mice (p=0.0128, unpaired t-test). **c** the pancreas showed no significant difference.

Error bars represent mean ± standard error of the mean (SEM). Significance level was set at *p* ≤ 0.05.*p≤0.05.

**Figure S2: Peripheral organ weight in ICV-treated WT and 3xTg-AD mice**

**a**-**c** Organ weight in grams (g), n=8 animals/group. **a** liver weight showed no significant treatment or strain changes. **b** spleen weight showed significant strain effect (F(1,29) =8,929, p<0.0001). Post hoc analysis showed a significant difference between vehicle WT-V/3xTg-V (p=0.0157) and SCDi treated WT-S/3xTg-S (p=0.0009) but no significant drug effect, 3xTg-V/3xTg-S (p=0.5725), 2-way ANOVA Tukey’s post hoc, **c** the pancreas showed no significant differences between any of the groups.

Error bars represent mean ± standard error of the mean (SEM). Significance level was set at *p* ≤ 0.05.**p≤0.01,****p≤0.0001.

## Notes

### Competing Interest Statement

The authors have declared no competing interest.

