## Supplementary figures and images for "Central inhibition of Stearoyl-CoA Desaturase has minimal effects on the peripheral metabolic symptoms of the 3xTg Alzheimer’s disease mouse model"

### Supplemental Figure 1

Supplemental Figure 1

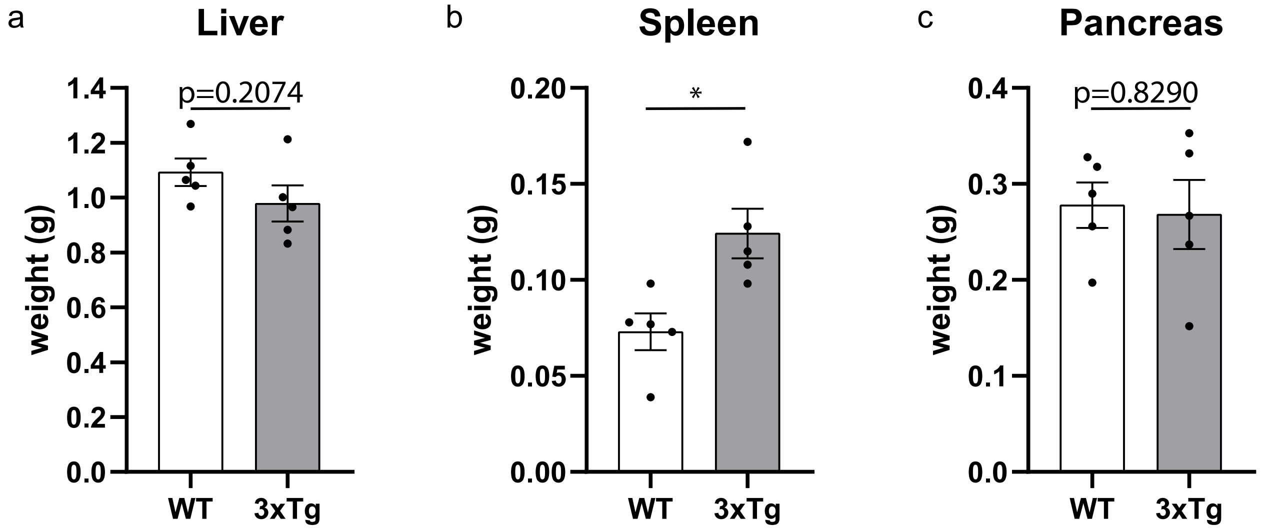

### Supplemental Figure 2

Supplemental Figure 2

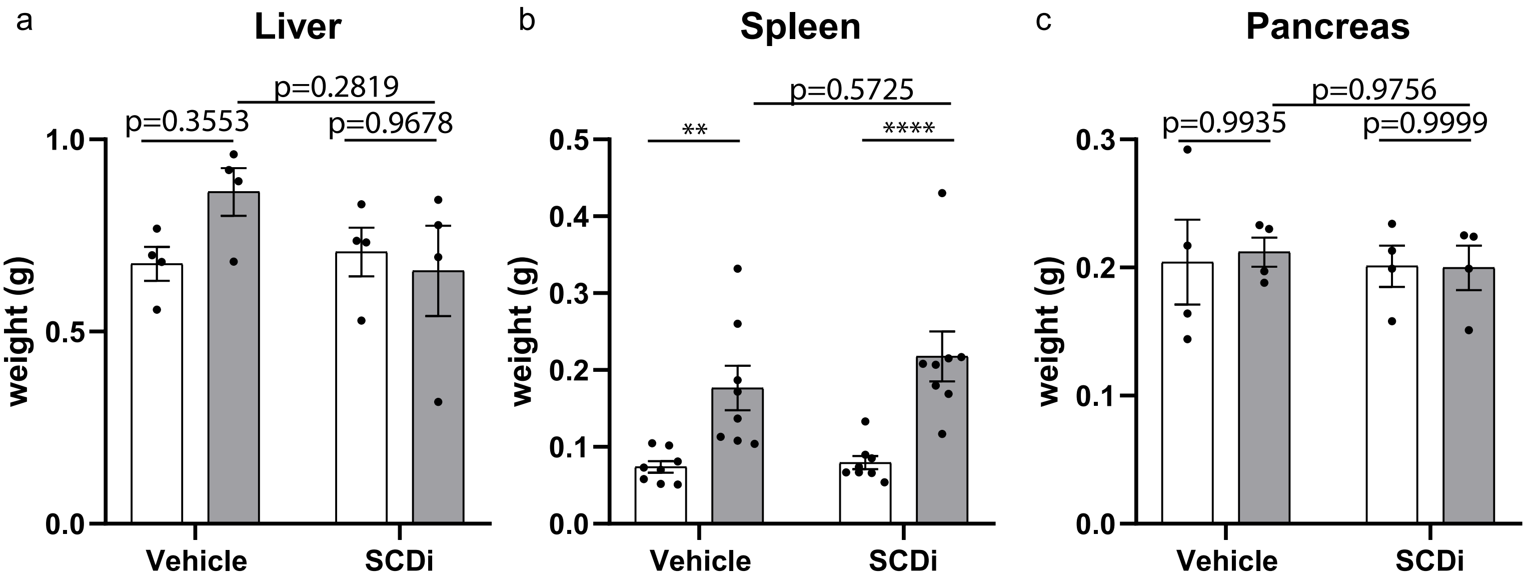
